# Intranasal fusion inhibitory lipopeptide prevents direct contact SARS-CoV-2 transmission in ferrets

**DOI:** 10.1101/2020.11.04.361154

**Authors:** Rory D. de Vries, Katharina S. Schmitz, Francesca T. Bovier, Danny Noack, Bart L. Haagmans, Sudipta Biswas, Barry Rockx, Samuel H. Gellman, Christopher A. Alabi, Rik L. de Swart, Anne Moscona, Matteo Porotto

**Affiliations:** Department Viroscience, Erasmus MC, Rotterdam, the Netherlands; Department of Pediatrics, Columbia University Medical Center, New York, NY, USA; Center for Host–Pathogen Interaction, Columbia University Medical Center, New York, NY, USA; Department of Experimental Medicine, University of Campania “Luigi Vanvitelli”, 81100 Caserta, Italy; Robert Frederick Smith School of Chemical and Biomolecular Engineering, Cornell University, Ithaca, New York, USA; Department of Chemistry, University of Wisconsin, Madison, WI, USA; Department of Microbiology & Immunology, Columbia University Medical Center, New York, NY, USA; Department of Physiology & Cellular Biophysics, Columbia University Medical Center, New York, NY, USA

**Author notes:** Authors contributed equally (Rory D. de Vries, Katharina S. Schmitz, Francesca T. Bovier). Contacts details: - Christopher A. Alabi - Rik L. de Swart - Anne Moscona - Matteo Porotto.

## Abstract

Containment of the COVID-19 pandemic requires reducing viral transmission. SARS-CoV-2 infection is initiated by membrane fusion between the viral and host cell membranes, mediated by the viral spike protein. We have designed a dimeric lipopeptide fusion inhibitor that blocks this critical first step of infection for emerging coronaviruses and document that it completely prevents SARS-CoV-2 infection in ferrets. Daily intranasal administration to ferrets completely prevented SARS-CoV-2 direct-contact transmission during 24-hour co-housing with infected animals, under stringent conditions that resulted in infection of 100% of untreated animals. These lipopeptides are highly stable and non-toxic and thus readily translate into a safe and effective intranasal prophylactic approach to reduce transmission of SARS-CoV-2.

**One-sentence summary:** A dimeric form of a SARS-CoV-2-derived lipopeptide is a potent inhibitor of fusion and infection *in vitro* and transmission *in vivo*.

## Main text

Infection by SARS-CoV-2 requires membrane fusion between the viral envelope and the host cell membrane, at either the cell surface or the endosomal membrane. The fusion process is mediated by the viral envelope spike glycoprotein, S. Upon viral attachment or uptake, host factors trigger large-scale conformational rearrangements in S, including a refolding step that leads directly to membrane fusion and viral entry (*1–6*). Peptides corresponding to the highly conserved heptad repeat (HR) domain at the C-terminus of the S protein (HRC peptides) may prevent this refolding and inhibit fusion, thereby preventing infection (*7–12*) (**Fig. 1a-c**). The HRC peptides form six-helix bundle (6HB)-like assemblies with the extended intermediate form of the S protein trimer, thereby disrupting the structural rearrangement of S that drives membrane fusion (*7*).

**Figure 1:**
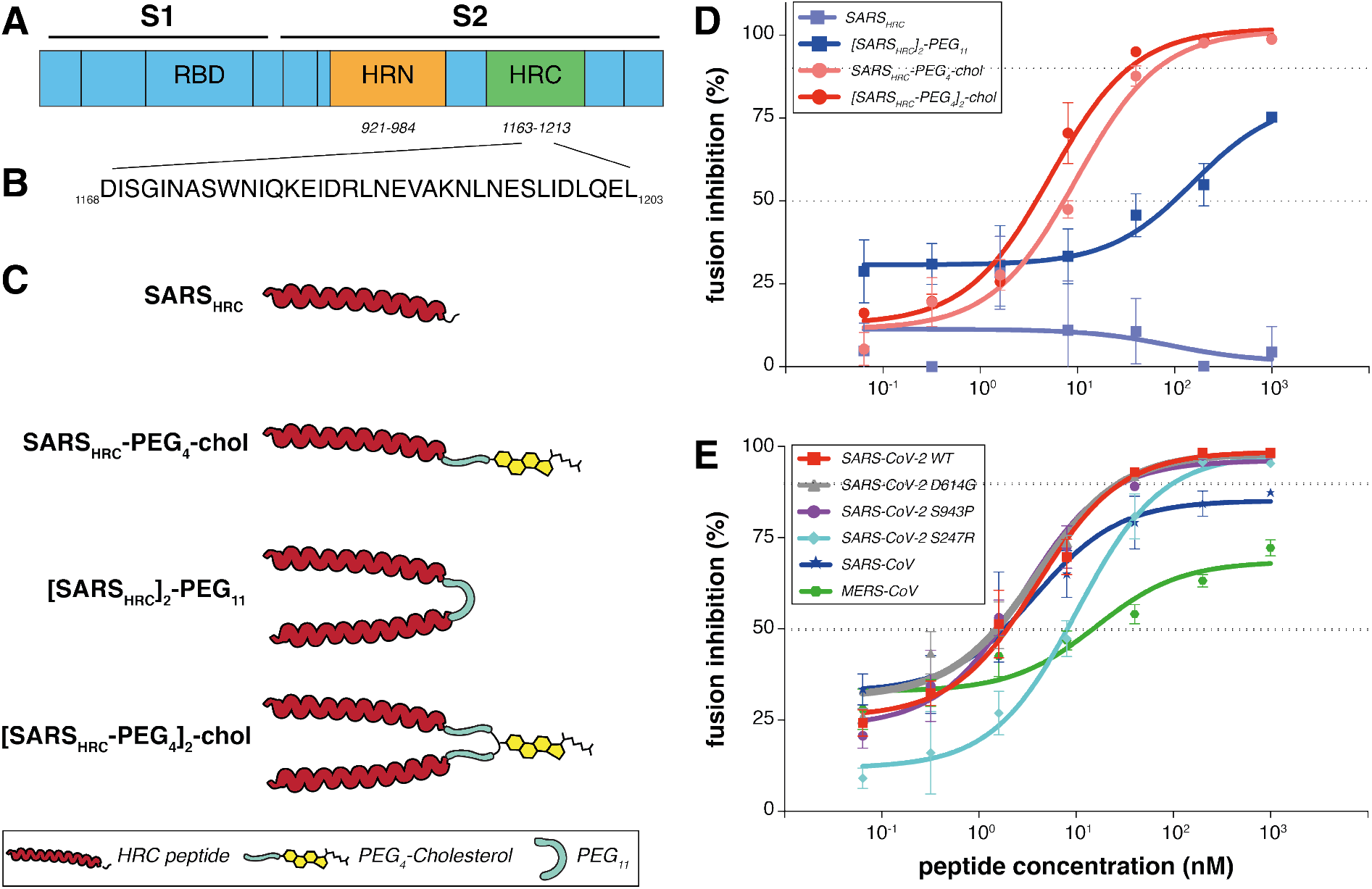
Peptide-lipid conjugates that inhibit SARS-CoV-2 spike (S)-mediated fusion. (A) The functional domains of SARS-CoV-2 S protein: receptor-binding domain (RBD) and heptad repeats (HRN and HRC) are indicated. (B) Sequence of the peptides that derive from the HRC domain of SARS-CoV-2 S. (C) Monomeric and dimeric forms of lipid tagged SARS-CoV-2 inhibitory peptides that were assessed in cell-cell fusion assays. (D) Cell-cell fusion assays with different inhibitory peptides. The percentage inhibition is shown for four different peptides used at increasing concentrations. Inhibitory concentrations 50% and 90% were calculated (dotted lines). Percent inhibition was calculated as the ratio of relative luminescence units in the presence of a specific concentration of inhibitor and the relative luminescence units in the absence of inhibitor and corrected for background luminescence. Data are means ± standard deviation (SD) from three separate experiments. The difference between the results for [SARS_HRC_-PEG_4_]_2_-chol and SARS_HRC_-PEG_4_-chol lipopeptides are significant (Two-way ANOVA, **\*\*\*\* p<0.0001**). (E) Fusion inhibitory activity of [SARS_HRC_-PEG_4_]_2_-chol peptide against SARS-CoV-2 S variants, MERS-CoV-2 S, and SARS-CoV S. Data are means ± standard deviation (SD) from three separate experiments.

We have previously demonstrated that lipid conjugation of HRC-derived inhibitory peptides markedly increases antiviral potency and *in vivo* half-life (*13–16*), and used this strategy to create entry inhibitors for prophylaxis and/or treatment of human parainfluenza virus type 3, measles virus, and Nipah virus infection (*14, 15, 17–20*). Both dimerization and peptide integration into cell membranes proved key to ensure respiratory tract protection and prevent systemic lipopeptide dissemination (*16, 18*). The lipid-conjugated peptides administered intranasally to animals reached high concentrations both in the upper and lower respiratory tract, and the specific nature of the lipid can be designed to modulate the extent of transit from the lung to the systemic circulation and organs (*18–22*). Lipid conjugation also enabled activity against viruses that do not fuse until they have been taken up via endocytosis (*23*). Here, we show that a dimeric form of a SARS-CoV-2 S-specific lipopeptide is a potent inhibitor of fusion mediated by the SARS-CoV-2 S protein, prevents viral entry, and, when administered intranasally, completely prevents direct-contact transmission of SARS-CoV-2 in ferrets. We propose this compound as a candidate antiviral, to be administered by inhalation or intranasal spray, for pre-exposure or early post-exposure prophylaxis for SARS-CoV-2 transmission in humans.

To improve the antiviral potency of the previously assessed SARS-CoV-2 HRC-lipopeptide fusion inhibitor (*7*), we compared monomeric and dimeric derivatives of the SARS-CoV-2 S-derived HRC-peptide (**Fig. 1d**). Initial functional evaluation of the SARS-CoV-2 HRC lipopeptides was conducted with a cell-cell fusion assay based on β-galactosidase (β-gal) complementation that we adapted for assessment of SARS-CoV-2 S-mediated fusion. For this assay, cells expressing human angiotensin-converting enzyme 2 (hACE2) and the N-terminal fragment of β-gal were mixed with cells expressing the SARS-CoV-2 S protein and the C-terminal fragment of β-gal. When fusion mediated by S occurs, the two fragments of β-gal combine to generate a catalytically active species, and fusion is detected via the luminescence that results from substrate processing by β-gal. This assay format allows the assessment of potential SARS-CoV-2 S-mediated membrane fusion inhibitors without the use of infectious virus. The assay measures an inhibitor’s ability to block fusion of S-bearing cells with receptor-bearing target cells and is predictive of *in vivo* antiviral activity (*14*).

**Fig. 1d** shows the antiviral potency of two monomeric and two dimeric SARS-CoV-2 S-derived 36-amino acid (**Fig. 1b**) HRC-peptides, without (SARS_HRC_ and [SARS_HRC_-PEG_4_]_2_) or with (SARS_HRC_-PEG_4_-chol and [SARS_HRC_-PEG_4_]_2_-chol) appended cholesterol, in cell-cell fusion assays. The percentage inhibition corresponds to the extent of luminescence signal suppression observed in the absence of any inhibitor (*i.e*., 0% inhibition corresponds to maximum luminescence signal). Dimerization increased the peptide potency (see SARS_HRC_ *vs*. [SARS_HRC_-PEG_4_]_2_ in **Fig. 1d**). The dimeric form of HRC lipopeptide was also more potent than its monomeric lipopeptide counterpart (SARS_HRC_-PEG_4_-chol IC_50_ ~10 nM and [SARS_HRC_-PEG_4_]_2_-chol IC_50_~3nM, (Two-way ANOVA, *p*<0.0001)). This dimeric cholesterol-conjugated peptide ([SARS_HRC_-PEG_4_]_2_-chol; red line in **Fig. 1d**) is the most potent lipopeptide against SARS-CoV-2 that has been identified thus far. A lipopeptide based on the human parainfluenza virus type 3 (HPIV3) F protein HRC domain, used as a negative control, did not inhibit fusion at any concentration tested (**Fig. S1**). A cellular toxicity (MTT) assay was performed in parallel with this experiment to evaluate the potential toxicity of each lipopeptide (**Fig. S2**). Toxicity for each of the lipopeptides in this assay was minimal, even at the highest concentrations tested (<20% at 100 μM). No toxicity was observed for the dimeric SARS-CoV-2 lipopeptide at its IC_90_ entry inhibitory concentration (~350 nM).

Despite the overall stability of the SARS-CoV-2 genome, variants with mutations in S have spread globally (*24–32*). These mutations in S altered infectivity of cells (*e.g*., D614G (*24*)) or were located in the putative target domain of the HRC peptide (*e.g*., S943P). To determine the potency of the [SARS_HRC_-PEG_4_]_2_-chol peptide for a range of variant SARS-CoV-2 viruses, we examined fusion inhibition mediated by each of these emerging S protein mutants. In addition, to assess the potential for broadspectrum activity we assessed potency against the S of SARS-CoV and MERS-CoV (using dipeptidyl peptidase 4 (DPP4) receptor-bearing cells as the target for the latter). **Fig. 1e** shows the IC_50_ and IC_90_ of [SARS_HRC_-PEG_4_]_2_-chol for inhibition of fusion by the S mutants and in addition, for SARS-CoV S and MERS-CoV S. The [SARS_HRC_-PEG_4_]_2_-chol lipopeptide inhibited all SARS-CoV-2 strains with S mutations at comparable potency and showed considerable potency against both SARS-CoV and MERS-CoV.

The lead peptide, [SARS_HRC_-PEG_4_]_2_-chol, was subsequently assessed for its ability to block entry of live SARS-CoV-2 in VeroE6 and VeroE6 cells overexpressing the protease TMPRSS2 (*33*), one of the host factors thought to facilitate viral entry at the cell membrane (*2*). The TMPRSS2-expressing cells accurately represent the entry route in airway cells, an important feature highlighted by the failure of chloroquine to inhibit infection in TMPRSS2-expressing Vero cells and in human lung cells (*34*). The [SARS_HRC_-PEG_4_]_2_-chol peptide was dissolved in an aqueous buffer containing 2% dimethylsulfoxide (DMSO), was incubated with cells for 1 hr at 37 °C, after which a fixed concentration of SARS-CoV-2 was added. After 8 hrs at 37 °C, fusion events were quantified by SARS-CoV-2 nucleoprotein (NP) staining. The [SARS_HRC_-PEG_4_]_2_-chol peptide inhibited live virus entry with an IC_50_ ~300 nM in cells not overexpressing TMPRSS and ~5 nM in VeroE6-TMPRSS2, with an IC_90_ ~1 uM in both cell types (**Fig. 2a**). In addition, we assessed the efficacy of the [SARS_HRC_-PEG_4_]_2_-chol peptide dissolved in a sucrose solution instead of DMSO, which would strengthen translational potential for human use. [SARS_HRC_-PEG_4_]_2_-chol peptide retained its potency in this formulation, with an IC_50_ ~300 nM in cells not overexpressing TMPRSS2 and ~5 nM in VeroE6-TMPRSS2 (**Fig. 2b**). The efficacy data are summarized in **Fig. 2c**.

**Figure 2.**
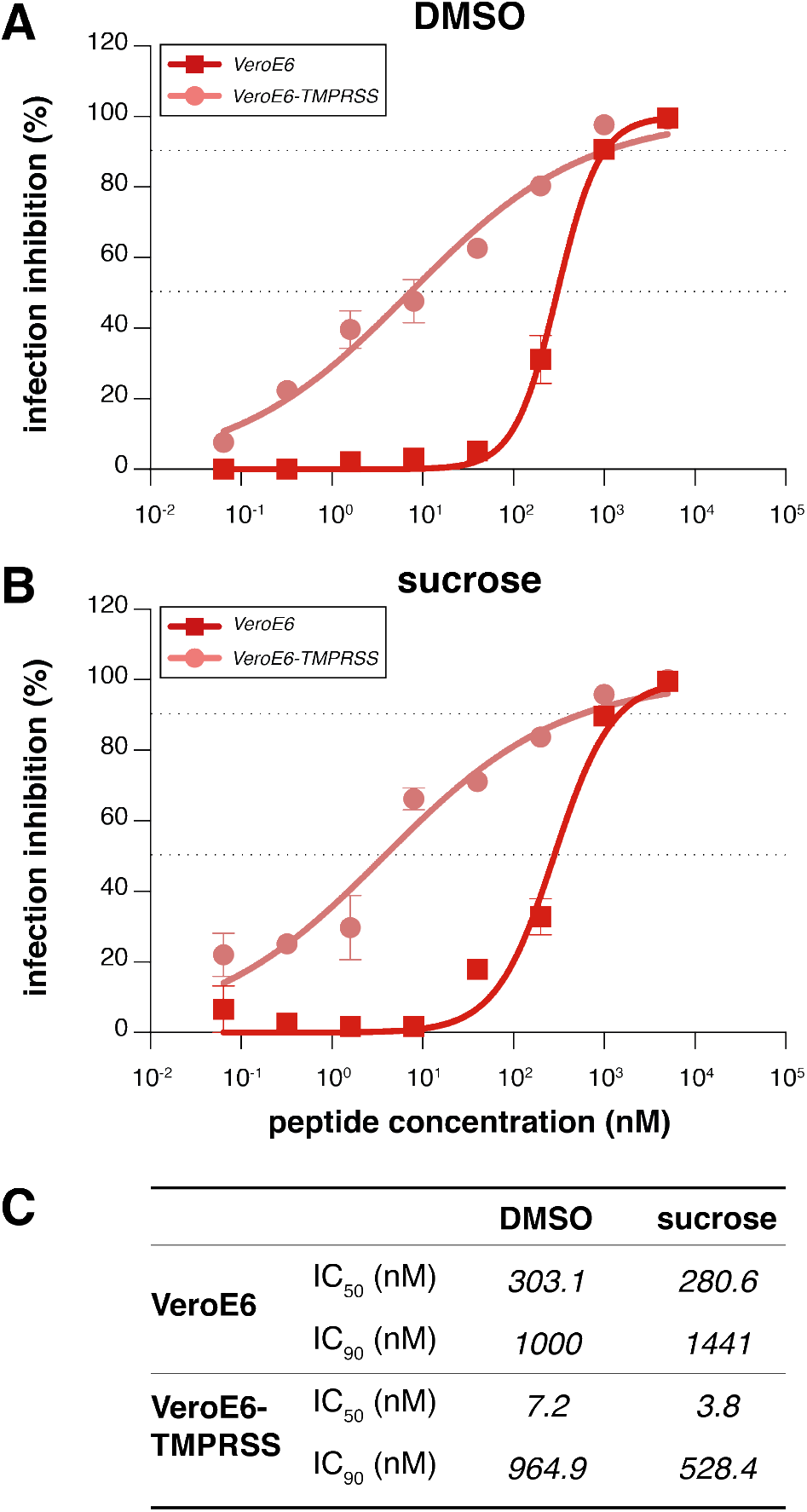
Inhibition of live SARS-CoV-2 entry by [SARS-CoV-2-HRC-peg_4_]_2_-chol peptide. The percentage inhibition of infection is shown on VeroE6 and VeroE6-TMPRSS2 cells with increasing concentrations of [SARS-CoV-2-HRC-peg_4_]_2_-chol. A DMSO-diluted stock (A, as used in ferrets) and sucrose-diluted stock (B, potential formulation for human use) were tested side-by-side. Inhibitory concentrations 50% and 90% were calculated (dotted lines). (C) Inhibitory concentrations 50% and 90% of [SARS_HRC_-PEG_4_]_2_-chol in live SARS-CoV-2 viral infection assays in VeroE6 cells with or without TMPRSS2 protease overexpression.

Ferrets are an ideal model for assessing respiratory virus transmission, either by direct contact or by aerosol transmission **(*35–40*)**. Mustelids are highly susceptible to infection with SARS-CoV-2, as also illustrated by frequent COVID-19 outbreaks at mink farms **(*41*).** Direct contact transmission of SARS-CoV in ferrets was demonstrated in 2003 **(*42*)**, and both direct contact and airborne transmission have recently been shown in ferrets for SARS-CoV-2 **(*36, 43, 44*)**. Direct contact transmission in the ferret model is highly reproducible (100% transmission from donor to acceptor animals), but ferrets display limited clinical signs. After infection via direct inoculation or transmission, SARS-CoV-2 can readily be detected in and isolated from the throat and nose, and viral replication leads to seroconversion.

To assess the efficacy of [SARS_HRC_-PEG_4_]_2_-chol in preventing SARS-CoV-2 transmission, naive ferrets were treated prophylactically with the lipopeptide before being co-housed with SARS-CoV-2 infected ferrets. In this setup, transmission via multiple routes can theoretically occur (aerosol, orofecal, and scratching or biting during play or fight), and ferrets are continuously exposed to infectious virus during the period of co-housing, providing a stringent test for antiviral efficacy. The study design is shown in **Fig. 3a**. Three donor ferrets (grey in diagram) were inoculated intranasally with 5.4 x 10^5^ TCID_50_ SARS-CoV-2 on day 0. Twelve recipient ferrets housed separately were treated by nose drops with a mock preparation (red) or [SARS_HRC_-PEG_4_]_2_-chol peptide (green) on 1-and 2-days post-inoculation (DPI) of the donor animals. The [SARS_HRC_-PEG_4_]_2_-chol peptides for intranasal administration were dissolved to a concentration of 6 mg/mL in an aqueous buffer containing 2% DMSO, administering a final dose of 2.7 mg/kg to ferrets (450 uL, equally divided over both nostrils). Six hours after the second treatment on 2 DPI, one infected donor ferret was co-housed with four naive recipient ferrets (two mock-treated, two peptide-treated). The experiment was performed in three separate, negatively pressurized HEPA-filtered ABSL3-isolator cages. After a 24-hour transmission period, co-housing was stopped and donor, mock-treated and peptide-treated ferrets were housed as separate groups. Additional [SARS_HRC_-PEG_4_]_2_-chol peptide treatments were given to recipient animals on 3 and 4 DPI. Peptide stocks and working dilutions had similar IC_50_ s, confirming that peptide-treated ferrets were always dosed with comparable amounts (**Fig. S3a and 3b)**.

**Figure 3.**
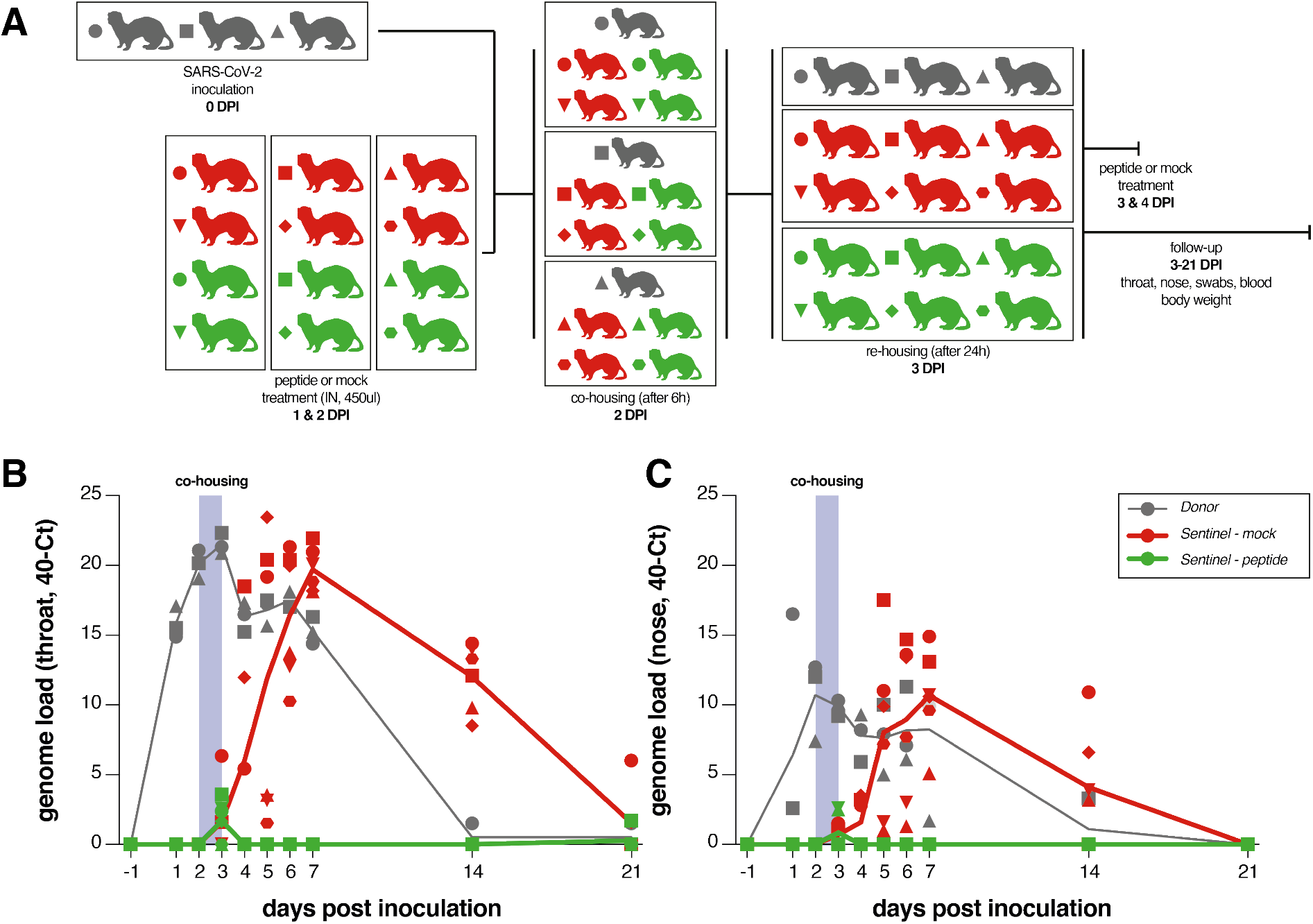
[SARS_HRC_-PEG_4_]_2_-chol prevents SARS-CoV-2 transmission *in vivo*. (a) Experimental design. (b) Viral loads detected in throat and nose swabs via RT-PCR. Viral loads are displayed as 40-Ct. Donor animals shown in grey, mock-treated animals in red, peptide-treated animals in green. 3/3 donor animals, 6/6 mock-treated animals and 0/6 lipopeptide-treated animals were productively infected. Symbols correspond to individual animals and are consistent throughout figures.

Throat and nose swabs were collected from ferrets daily for the first week, and additionally on 14 and 21 DPI, for assessment of viral replication. Small volume blood samples were collected on 0, 7, 14, and 21 DPI for assessing the presence of neutralizing antibodies in serum. **Fig. 3b and 3c** shows the viral loads (detection of viral genomes via RT-qPCR) for directly inoculated donor animals (grey), mock-treated recipient animals (red) and lipopeptide-treated recipient animals (green). All directly inoculated donor ferrets were productively infected, as shown by SARS-CoV-2 genome detection in throat and nose swabs, and efficiently and reproducibly transmitted the virus to all mock-treated acceptor ferrets (**Fig. 3b and 3c**, red curves). Notably, productive SARS-CoV-2 infection was not detected in the throat or nose of any of the peptide-treated recipient animals (**Fig. 3b and 3c**, green curves). A slight rise in viral loads in samples collected at 3DPI was detected, at the end of the co-housing, confirming that peptide-treated animals were exposed to SARS-CoV-2. Strikingly, from 4 DPI onwards, there was a clear treatment effect in which the [SARS_HRC_-PEG_4_]_2_-chol peptide protected 6/6 ferrets from transmission and productive infection. Donor ferrets and 6/6 mock-treated recipient animals seroconverted on 21 DPI. None of the peptide-treated animals seroconverted, demonstrating that in-host virus replication was completely blocked by [SARS_HRC_-PEG_4_]_2_-chol*J* (**Fig. 4**). None of the animals showed clinical signs as a result of infection or treatment over the course of the experiment, and body weights remained stable (**Fig. S4**).

**Figure 4.**
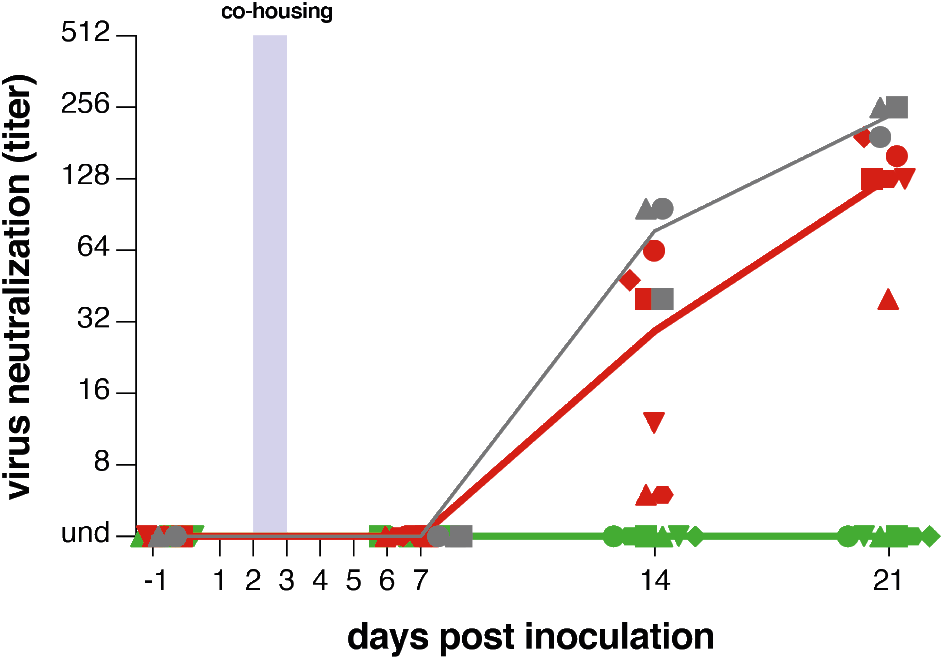
[SARS_HRC_-PEG_4_]_2_-chol-treated animals do not seroconvert. Presence of neutralizing antibodies was determined in a live virus neutralization assay. Virus neutralizing antibodies are displayed as endpoint serum dilution factor to block SARS-CoV-2 replication. Donor animals shown in grey, mock-treated animals in red, peptide-treated animals in green. 3/3 donor animals, 6/6 mock-treated animals and 0/6 lipopeptide-treated animals seroconverted. Symbols correspond to individual animals and are consistent throughout figures.

Based on the *in vitro* and *in vivo* results shown here, we expect that prophylactic intranasal administration of the [SARS_HRC_-PEG_4_]_2_-chol peptide prevents transmission from infected to uninfected individuals, even during a 24-hour period of intense direct contact. *In vitro* data suggest that this lipopeptide will be effective against emerging variants with mutations in S and possibly against other coronaviruses. This efficacy can be readily assessed in real time and adjustments made if needed.

Parallel approaches to prevent transmission that target ACE2 or the interaction between S and ACE2 have also shown promise *in vitro* (e.g. the “miniprotein” approach recently reported by Cao *et al* (*45*)). In distinction to various antibody or nanobody products (*46*) the [SARS_HRC_-PEG_4_]_2_-chol peptide is inexpensive to produce, has a long shelf life, and does not require refrigeration. Moreover, this is the first compound to convincingly prevent SARS-CoV-2 transmission in a relevant animal model. We envision the use of fusion inhibitory lipopeptides as complementary to other pandemic mitigation strategies. In addition to the nasal drop administration for the [SARS_HRC_-PEG_4_]_2_-chol peptide, other routes that would be equally or more acceptable to humans, for example inhalation devices, are being explored. This HRC lipopeptide fusion inhibitor is feasible for advancement to human use and should readily translate into a safe and effective nasal spray or inhalation administered fusion inhibitor for anti-SARS-CoV-2 prophylaxis, thus supporting containment of the current COVID-19 pandemic.

## Acknowledgements

We thank Mart Lamers, Sander Herfst, Elwin Verveer, Anna Mykytyn and Marion Koopmans for their contributions to this study.

## Author contributions

conceptualization RDdV, SHG, CAA, RLdS, AM, MP; formal analysis RDdV, FTB, CAA, KSS, MP; funding acquisition BLH, RLdS, AM, MP; investigation RDdV, FTB, KSS, DN, SHG, CAA, RLdS, MP; resources BLH, BR, CAA, RLdS, AM, MP; supervision RLdS, CAA, AM, MP; visualization RDdV, KS, AM, MP; writing – original draft RDdV. RLdS, AM, MP; final version: all coauthors provided feedback to the final draft.

## Data and materials availability

All data is available in the manuscript or the supplementary materials.

## Supplementary Materials

### List of Supplementary Materials

Materials and Methods

Figure S1 – S5

### Materials and Methods

#### Ethics statement

Influenza virus, SARS-CoV-2 and Aleutian Disease Virus seronegative female ferrets (*Mustela putorius furo*), weighing 900-1200g, were obtained from a commercial breeder (Triple F Farms, PA, USA). Animals were housed and experiments were performed in compliance with the Dutch legislation for the protection of animals used for scientific purposes (2014, implementing EU Directive 2010/63). Research was conducted under a project license from the Dutch competent authority (license number AVD1010020174312) and the study protocol was approved by the institutional Animal Welfare Body (Erasmus MC permit number 17-4312-07). Animal welfare was monitored on a daily basis.

#### Lipopeptide synthesis

The peptide (SARS_HRC_) corresponding to residues 1168–1203 of SARS-CoV-2 S with a C-terminal -GSGSGC spacer sequence was prepared by solid phase peptide synthesis (SPPS). The SARS_HRC_ peptide was acetylated at the N-terminus and amidated at the C-terminus. The crude peptide was purified by reversed-phase HPLC chromatography and characterization by MALDI-TOF MS. The SARS_HRC_-PEG_4_-chol, [SARS_HRC_]_2_-PEG_11_, and [SARS_HRC_-PEG_4_]_2_-chol were synthesize d via chemoselective thiol-Michael addition reactions between the terminal thiol group on the peptide cysteine residue and maleimide functional PEG linkers or PEG-cholesterol linkers as previously described (*1*). HPLC purification and lyophilization yielded the peptide-lipid conjugates as white powders. The identity of the conjugates was verified by MALDI-TOF MS (**Fig. S5).**

#### Dissolving [SARS_HRC_-PEG_4_]_2_-chol for use in experiments

[SARS_HRC_-PEG_4_]_2_-chol was supplied as a white powder in aliquots of 10 mg. For *in vitro* experiments with live virus and *in vivo* experiments in ferrets, 10 mg of [SARS_HRC_-PEG_4_]_2_-chol was dissolved in 33.3 ul DMSO, which was subsequently added to 1632.7 ul de-ionized H_2_O. This yielded a final aqueous solution of lipopeptide dissolved at a concentration of 6 mg/mL containing 2% DMSO. To obtain peptide dissolved in aqueous solution without DMSO, 100 mg/ml of the [SARS_HRC_-PEG_4_]_2_-chol peptide in DMS0 (10 mg of peptide in 100 ul of DMSO) and 1 mg/ml of sucrose in sterile water were prepared. 10 ul of the peptide solution (1mg) was added to 100μl of sucrose (0.1 mg). Lyophilisation of the peptide solution (DMSO + sucrose) was performed over-night and dry powder was resuspended in 50μl of sterile water to a final concentration is 20 mg/ml in water without any DMSO.

#### Cells

Human kidney Epithelial (HEK) 293T and Vero (African green monkey kidney) cells were grown in Dulbecco’s modified Eagle’s medium (DMEM; Invitrogen; Thermo Fisher Scientific) supplemented with 10% foetal bovine serum (FBS) and antibiotics in 5% CO_2_ at 37°C. VeroE6 (ATCC CRL-1586) and VeroE6-TMPRSS2 cells were grown in DMEM (Gibco) with 10% FBS, 2 mM L-glutamine (Gibco), 10 mM Hepes (Lonza), 1.5 mg/ml sodium bicarbonate (NaHCO3, Lonza), penicillin (10,000 IU/mL) and streptomycin (10,000 IU/mL)(*2*).

#### HAE cultures

The EpiAirway AIR-100 system (MatTek Corporation) consists of normal human-derived tracheal/bronchial epithelial cells that have been cultured to form a pseudostratified, highly differentiated mucociliary epithelium closely resembling that of epithelial tissue *in vivo*. Cultures were transferred to six-well plates containing 1.0 ml medium per well (basolateral feeding, with the apical surface remaining exposed to air) and acclimated at 37°C in 5% CO2 for 24h prior to experimentation (*3*).

#### Plasmids

The cDNAs coding for hACE2 fused to the fluorescent protein Venus, dipeptidyl peptidase 4 (DPP4) fused to the fluorescent protein Venus, SARS-CoV-2 S, SARS-CoV S, and MERS-S (codon optimized for mammalian expression) were cloned in a modified version of the pCAGGS (with puromycin resistance for selection).

#### β-Gal complementation-based fusion assay

We previously adapted a fusion assay based on alpha complementation of β-galactosidase (β-Gal)(*4*). In this assay, hACE2 receptor-bearing cells (or dipeptidyl peptidase 4 (DPP4) receptor-bearing cells for MERS-CoV-2 experiments) expressing the omega peptide of β-Gal are mixed with cells co-expressing glycoprotein S and the alpha peptide of β-Gal, and cell fusion leads to alpha-omega complementation. Fusion is stopped by lysing the cells and, after addition of the substrate (^®^The Tropix Galacto-Star™ chemiluminescent reporter assay system, Applied Biosystem), luminescence is quantified on a Tecan M1000PRO microplate reader.

#### Cell toxicity assay

HEK293T or Vero cells were incubated with the indicated concentration of the peptides or vehicle (dimethyl sulfoxide) at 37 °C. Cell viability was determined after 24h using the Vybrant MTT Cell proliferation Assay Kit according to the manufacturer’s guidelines. The absorbance was read at 540 nm using Tecan M1000PRO microplate reader. HAE cultures were incubated at 37°C in the presence or absence of 1, 10, or 100 μM concentrations of the peptide, and peptide was added to the feeding medium every 2 days for 7 days. Cell viability was determined on day 7 as above.

#### Virus

SARS-CoV-2 (isolate BetaCoV/Munich/BavPat1/2020; kindly provided by Prof. Dr. C. Drosten) was propagated to passage 3 on VeroE6 cells in OptiMEM I (1X) + GlutaMAX (Gibco), supplemented with penicillin (10,000 IU/mL, Lonza) and streptomycin (10,000 IU/mL, Lonza) at 37°C. VeroE6 cells were inoculated at a multiplicity of infection (MOI) of 0.01. Supernatant was harvested 72 hours post inoculation (HPI), cleared by centrifugation and stored at −80°C. All live virus work was performed in a Class II Biosafety Cabinet under BSL-3 conditions at Erasmus MC.

#### *In vitro* potency of HRC dimer-chol

Potency of [SARS_HRC_-PEG_4_]_2_-chol was determined in an *in vitro* live virus fusion assay. Original stocks and working dilutions for animal experiments were tested in triplicate in VeroE6 and VeroE6-TMPRSS2 cells at concentrations of 0.06 nM to 5 μM. Peptide was pre-incubated with the cells for 1 hr at 37°C. After pre-incubation, virus (600 TCID_50_) was added. After 8 hrs at 37°C, cells were washed and fixed with 4% PFA for 20 min at room temperature (RT). Plates were submerged in 70% ethanol and stained in a BSL-2 laboratory. In short, cells were washed with PBS and blocked with 10% normal goat serum (NGS) for 30 min at RT. Primary mouse-anti-SARS-CoV nucleocapsid antibody (1:1000, Biorad) was incubated for 1 hr at RT in 10% normal goat serum (NGS). After washing, secondary goat-anti-mouse IgG/FITC antibody (1:1000, Invitrogen) was incubated for 45 min at RT in 10% NGS. Fluorescent spots were visualized with an Amersham Typhoon Biomolecular Imager (GE Healthcare) and counted with ImageQuant TL 7.0 software (GE Healthcare).

#### Ferret transmission experiment

All animal handlings were performed under anaesthesia with a mixture of ketamine/medetomidine (10mg/kg and 0.05mg/kg, respectively) antagonized by atipamezole (0.25 mg/kg). Three donor ferrets were inoculated intranasally with 5.4 x 10^5^ TCID_50_ of SARS-CoV-2 in 450μl (225μl instilled dropwise in each nostril) and were housed together in a negatively pressurized HEPA-filtered ABSL-3 isolator. This was considered the start of the experiment (0 days post inoculation, DPI). At the same time, twelve direct contact ferrets were divided over three other isolators. Ferrets were either mock-treated (vehicle, 2%DMSO in distilled water) or treated with [SARS_HRC_-PEG_4_]_2_-chol on 1-4 DPI. The peptide was inoculated intranasally in 450μl (225μl instilled dropwise in each nostril), HRC dimer-chol treated ferrets received a peptide dose of ~2.7 mg/kg. Leftover batches were stored at −80°C for later use in *in vitro* potency assays. At 2 DPI, six hours after the second treatment, one donor ferret was placed in the same isolators as two mock-treated and two peptide-treated ferrets, in three separate isolators (**Fig. 3a**). Each isolator now contained five ferrets, the donor ferret, the mock-treated recipient ferrets and the [SARS_HRC_-PEG_4_]_2_-chol-treated recipient ferrets. At 3 DPI, 18 hours after onset of cohousing, the animals received a third mock or peptide treatment, Six hours later, i.e. 24 hours after the start of the co-housing, the donor animals were moved back to their original isolator and the mock-treated and peptide-treated ferrets were housed in two groups of six animals in clean isolators.

Throat and nose swabs were collected from the animals on 0, 1, 2, 3, 4, 5, 6, 7, 14 and 21 DPI. Samples were always obtained prior to dosing with mock or peptide. Swabs were stored at −80°C in virus transport medium (Minimum Essential Medium Eagle with Hank’s BSS (Lonza), 5 g L-1 lactalbumine enzymatic hydrolysate, 10% glycerol (Sigma-Aldrich), 200 U/ml of penicillin, 200 mg/ml of streptomycin, 100 U/ml of polymyxin B sulfate (Sigma-Aldrich), and 250 mg/ml of gentamicin (Life Technologies)). Blood samples were obtained from ferrets on 0, 7, 14 and 21 DPI by vena cava puncture. Blood was collected in serum-separating tubes (Greiner), processed, heat-inactivated and sera were stored at −80°C. The study was stopped at 21 DPI. All animal experiments were performed in class III isolators in a negatively pressurized ABSL3 facility, all handlings were performed under general anaesthesia.

#### RNA isolation and RT-qPCR on throat and nose swabs

Sixty ul sample (virus transport medium in which swabs are stored) was added to 90 ul of MagNA Pure 96 External Lysis Buffer (Roche). A known concentration of phocine distemper virus (PDV) was added to the sample as internal control for the RNA extraction (*5*). The 150 ul of sample/lysis buffer was added to a well of a 96-well plate containing 50 ul of magnetic beads (AMPure XP, Beckman Coulter). After thorough mixing, the plate was incubated for 15 min at room temperature. The plate was then placed on a magnetic block (DynaMag™-96 Side Skirted Magnet (ThermoFisher Scientific)) and incubated for 3 min to allow the displacement of the beads towards the side of the magnet. Supernatants were carefully removed and beads were washed three times for 30 sec at room temperature with 200 ul/well of 70% ethanol. After the last wash, a 20 ul multichannel pipet was used to remove residual ethanol. Plates were air-dried for 2 min at room temperature. Plates were removed from the magnetic block and 50ul of PCR grade water was added to each well and mixed. Plates were incubated for 5 min at room temperature and then placed back on the magnetic block for 2 min to allow separation of the beads. Supernatants were pipetted in a new plate and RNA was stored at −80°C. The RNA was directly used for RTqPCR using primers and probes targeting the E gene of SARS-CoV-2 as previously described (*6*).

#### Virus neutralization of ferret sera

Seroconversion of ferrets was tested with ferret sera from 0, 7, 14 and 21 DPI. Duplicates of ferret sera were incubated with 100 TCID_50_ of virus in a 2-fold dilution series starting at a concentration of 1:8 for 1 hr at 37°C. Virus-sera mix was added to VeroE6 cells and incubated for 5 days at 37°C. Cytopathic effect was used as readout to determine the minimal serum concentration required to inhibit CPE formation.

**Figure S1.**
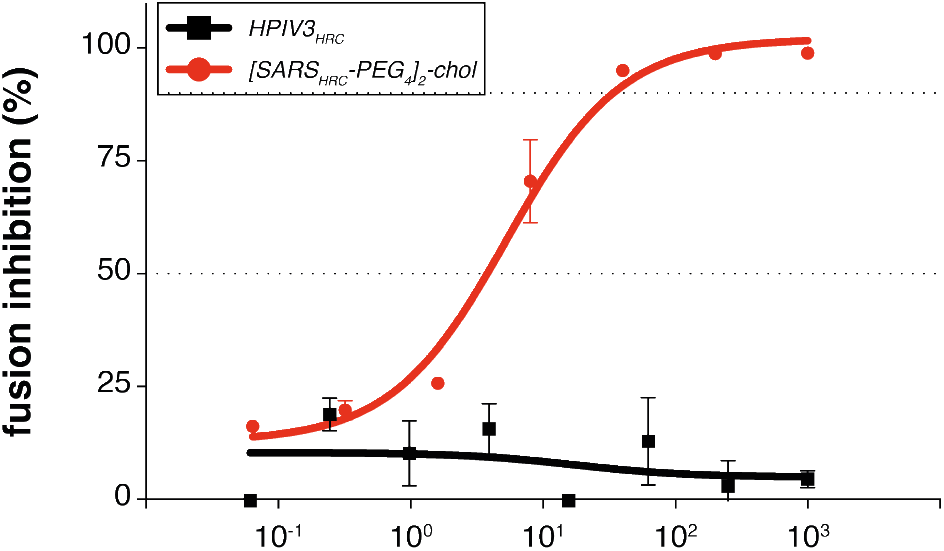
Specificity of SARS-CoV-2 inhibition by [SARS-CoV-2-HRC-peg_4_]_2_-chol. A lipopeptide based on the human parainfluenza virus type 3 (HPIV3) F protein HRC domain, used as a negative control, did not inhibit fusion at any concentration tested.

**Figure S2.**
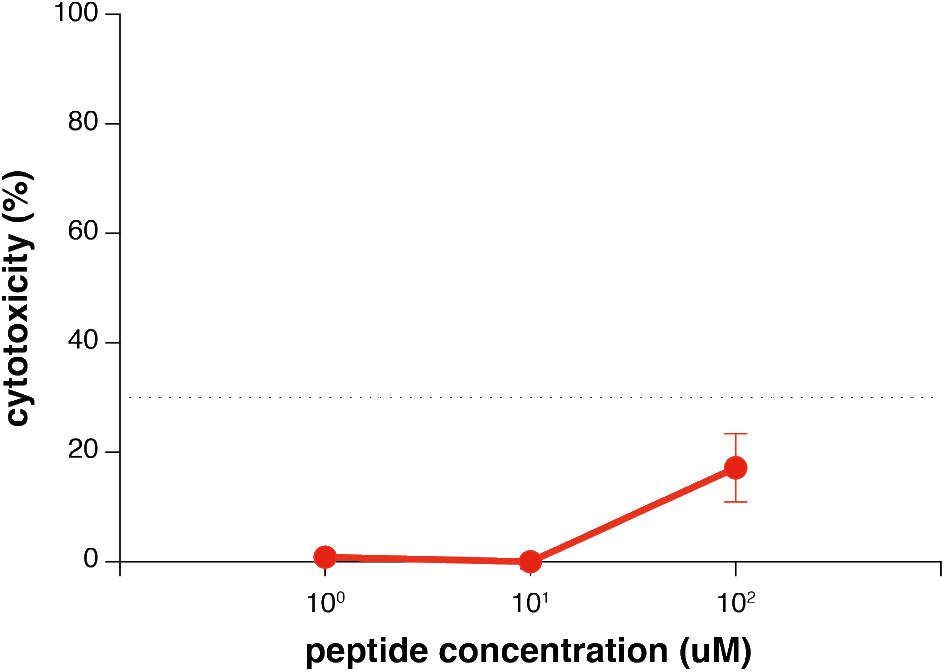
*Ex vivo* cytoxicity assessment. An MTT assay was used to determine the toxicity of the [SARS-CoV-2-HRC-peg_4_]_2_-chol in human airway epithelium (HAE). No toxicity was observed for the peptide at the concentrations of 1 and 10 μM. Toxicity was minimal (<20%) at the highest concentration tested (100 μM).

**Figure S3:**
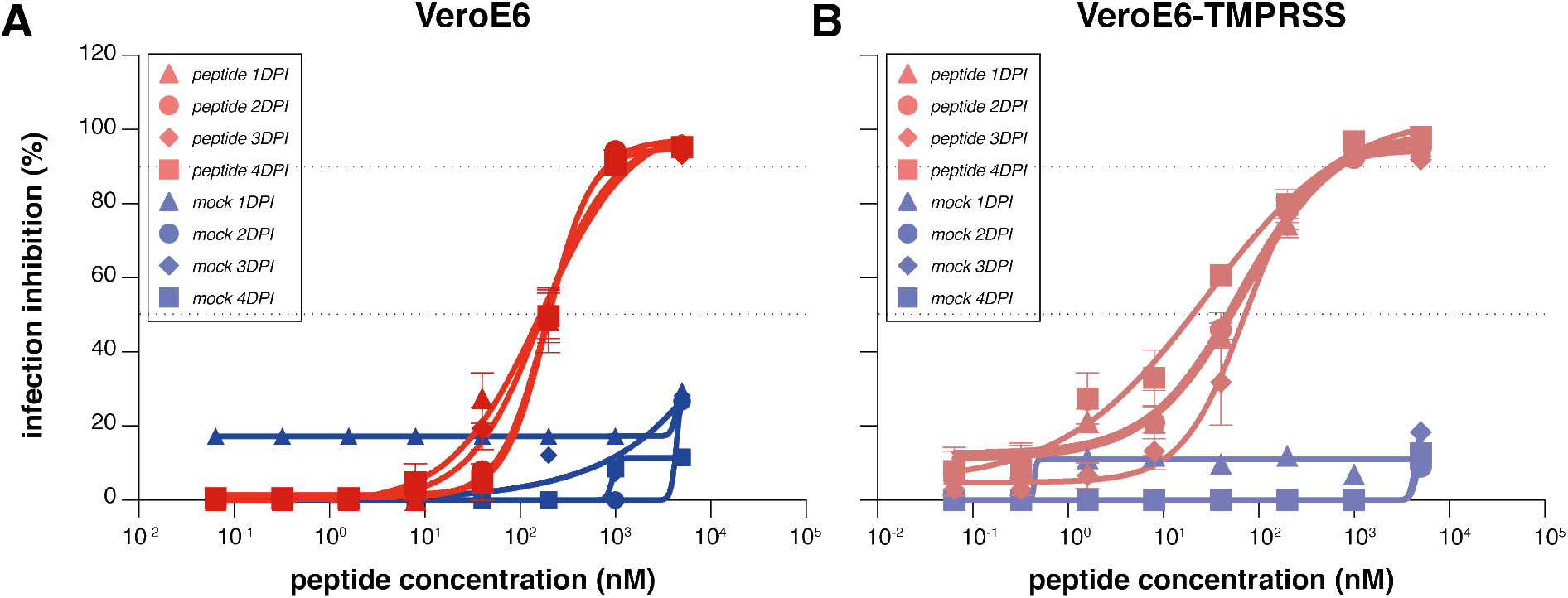
Potency of peptide stocks used *in vivo*. The potency of peptide dilutions used on 1-4 DPI in the *in vivo* experiments was confirmed with a live virus infection assay. The percentage infection events is shown on (A) VeroE6 and (B) VeroE6-TMPRSS with increasing concentrations of [SARS-CoV-2-HRC-peg_4_]_2_-chol (red) or mock (blue). Inhibitory concentrations 50% and 90% are indicated (dotted lines). Data are means ± standard error of the mean (SEM) from one experiment.

**Figure S4.**
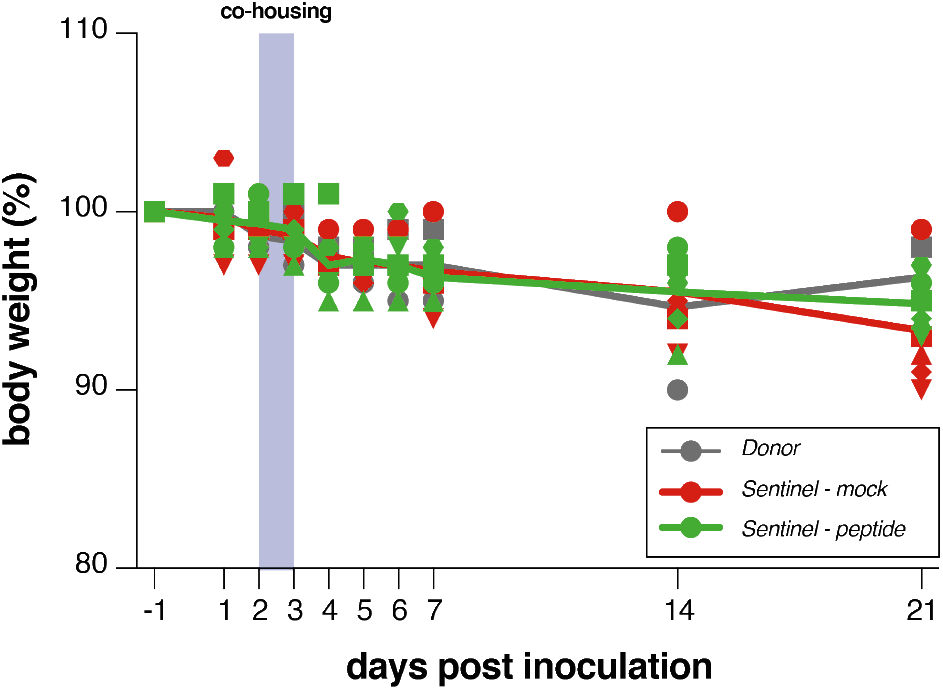
SARS-CoV-2-infected ferrets do not lose weight. Body weights of all ferrets remained stable over the time of the experiment. donor animals shown in grey, mock-treated animals in red, peptide-treated animals in green. Symbols correspond to individual animals and are consistent throughout figures.

**Figure S5.**
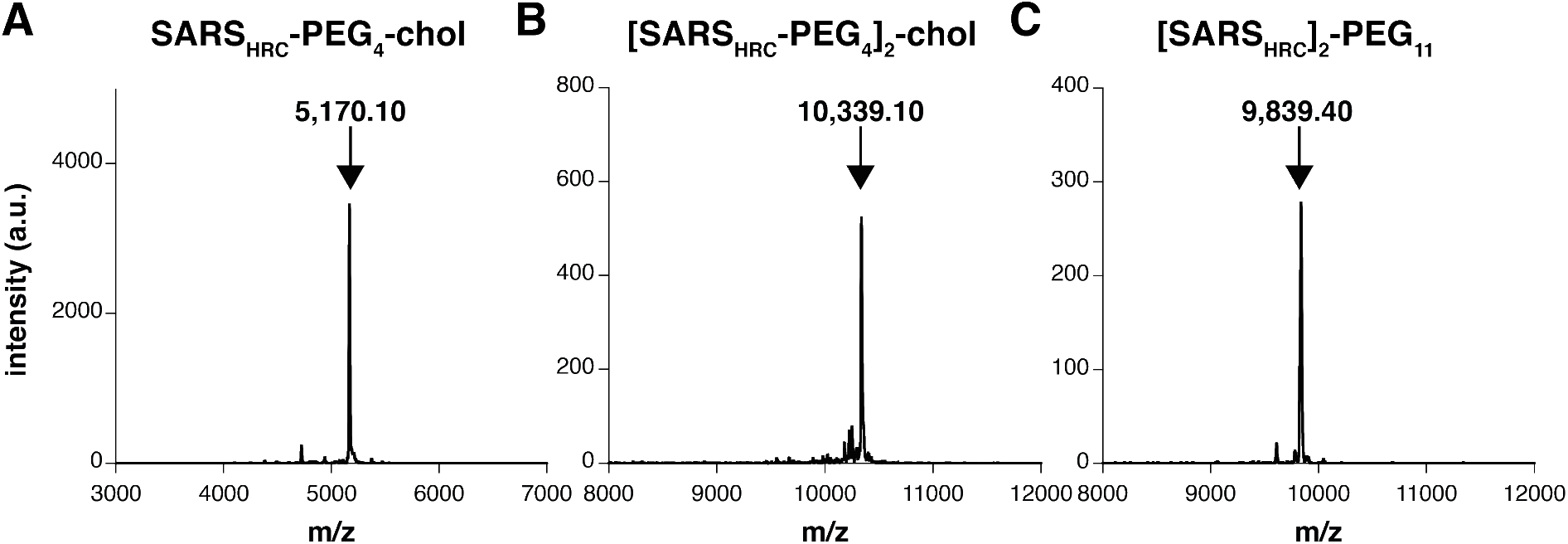
Identity of the conjugates was verified by MALDI-TOF MS. (A) MALDI of SARS_HRC_-PEG_4_-chol. Theoretical: 5170.8 Da; observed 5170.1 Da. (B) MALDI of [SARS_HRC_-PEG_4_]_2_-chol. Theoretical m/z: 10,335.4 Da; observed 10,339.10 Da. (C) MALDI of [SARS_HRC_]-PEG_11_. Theoretical m/z: 9841.0 Da; observed m/z: 9,839.40 Da.

